# Fus3 interacts with Gal83, revealing the MAPK crosstalk to Snf1/AMPK to regulate secondary metabolic substrates in filamentous fungi

**DOI:** 10.1101/2023.07.05.547858

**Authors:** Longxue Ma, Fuguo Xing, Xu Li, Bowen Tai, Ling Guo

**Affiliations:** Institute of Food Science and Technology, Chinese Academy of Agricultural Sciences, Ministry of Agriculture, Beijing 100193, P. R. China; Key Laboratory of Agro-products Quality and Safety Control in Storage and Transport Process, Ministry of Agriculture, Beijing 100193, P. R. China; Institute of Food Science and Technology, Chinese Academy of Agricultural Sciences, 2 Yuanmingyuan West Road, Haidian District, Beijing 100193, P. R. China

## Abstract

The pheromone MAPK is essential for the vital activities of fungi and is widely identified in filamentous fungi of agricultural, medical, and industrial relevance. The targets have rarely been reported and it is difficult to understand the mechanism of pheromone MAPK signaling pathway. Aflatoxins (AFs), highly carcinogenic natural products, are produced by the secondary metabolism of fungi, such as *Aspergillus flavus*. Our previous studies demonstrated that Fus3 regulates AFs by modulating substrate levels in *Aspergillus flavus*, but no mechanism explain that in fungi. Here we show Gal83, a new target of Fus3, and identified the pheromone Fus3-MAPK signaling pathway regulates the Snf1/AMPK energy-sensing pathway to modulate aflatoxins synthesis substrates. In the screening for target proteins of Fus3, the Snf1/AMPK complexes β subunit was identified by using tandem affinity purification and multi-omics, which physically interacted with Fus3 in *vivo* and *vitro* and received phosphorylation from Fus3. While neither aflatoxin transcript levels were down-regulated in *gal83*-mutant and *fus3*-mutant strains, significant decreases in aflatoxin B_1_, aflatoxin synthetic substrates levels and gene expression levels of primary metabolic enzymes were shown that both the Fus3-MAPK and Snf1/AMPK pathways could response energy signal. In conclusion, all the evidence unlocks a novel pathway of Fus3-MAPK to regulate AFs synthesis substrates by cross-talking to the Snf1/AMPK complexes.

**Importance:** Aflatoxin poses a great threat to human and animal health and the economy, thus the mechanisms regulating aflatoxin synthesis have been of great interest. We have previously demonstrated that MAPK regulates aflatoxin biosynthesis significantly, but the regulatory mechanism of Fus3-MAPK is not clear. Here we found that Pheromone Fus3-MAPK responds to energy and transmits to Snf1/AMPK through phosphorylation, which regulates the level of secondary metabolic substrates in *Aspergillus flavus*, as a novel pathway of Fus3-MAPK. Fus3 interacts stably with Gal83 and colocalizes in the cytoplasm and nucleus, directly regulating the levels of aflatoxin synthetic substrates. These data advance our understanding of the regulation of aflatoxin by pheromone MAPK, and the mechanism of pheromone MAPK and Snf1/AMPK crosstalk regulation is confirmed. Overall, this has a positive effect on both fungal regulatory mechanisms and aflatoxin prevention and control.

## Introduction

Mitogen-activated protein kinase (MAPK) pathways, which obtain information from G protein-coupled receptors (GPCR), are evolutionarily conserved in eukaryotic organisms from yeast to mammals, consist of three cascade proteins, MAPKKK, MAPKK and MAPK (1–5). The pheromone Fus3-MAPK pathway was first identified as an important signaling pathway for mating in yeast (6, 7). Although conserved, which regulates a wider range of functions in filamentous fungi, including growth rate, sexual development, pathogenicity, and secondary metabolism (8–13). Similar to yeast, in filamentous fungi, such as *Aspergillus nidulans*, *Aspergillus oryzae* and *Aspergillus flavus*, the Fus3 kinase, as the terminal key kinase protein of the Fus3-MAPK pathway, is thought to regulate fungi development and secondary metabolism by regulating transcription factor SteA/Ste12 in the nucleus and phosphorylate the global regulator VeA (1, 9, 14). In *Aspergillus oryzae*, AoFisA, presumed to be a co-interacting protein, could interact with AoFus3 and AoSte12 (15).

*Aspergillus flavus,* one of the main producers of the highly carcinogenic aflatoxins (AFs), is a filamentous pathogen of being high concern with the widespread infestation of agricultural crops and sideline products (16). AFs, being ingested at low doses over a long term may lead to immunosuppression, liver cancer and even death, which poses a great threat to humans and animals (17, 18). To reduce AFs biosynthesis at source, there is an urgent need to elucidate the regulating mechanisms. The previous studies have shown that Fus3 is essential for fungal development and secondary metabolism (1, 9–11). Interestingly, our previous study shown that Fus3 regulates AFs synthesis substrate levels rather than directly regulating AFs synthesis genes (1), and the existing regulatory mechanisms of Fus3 cannot explain this phenomenon.

In this work, to unravel the mechanisms of Fus3 regulation, particularly AFs, we focused on proteins with post-translational modifications and interaction conditions, identified the Gal83 was a new stable target for Fus3. We screened 133 proteins using a combination of tandem affinity purification techniques and phosphoprotein downregulation groups, and transcriptional level non-downregulation group. GO and KEGG enrichment analyses revealed that the screening proteins were closely associated with the use of carbon source selection. Only Gal83, the β subunit of Snf1/AMPK complexes was identified as receptor of Fus3. Importantly, the function of Gal83 was consistent with Fus3 by phenotype identification and transcriptome analysis, and importantly both regulated AFs substrate levels rather than directly regulating AFs gene clusters. Finally, we shown that Fus3-MAPK cross-talked to Snf1/AMPK to regulate the primary metabolism and the level of AFs synthesis substrate in *A. flavus*, which provides a new pathway model for fungal regulatory networks.

## RESULTS

### The ability to adapt and use carbon sources is lost in Δ*fus3* strain

The Fus3 regulates many mechanisms has been reported (1, 2, 15, 19), yet those are unclear in aflatoxins (AFs) synthesis. Significantly different phenotypes of the Δ*fus3* strain in different carbon sources, which were researched carefully, such as PDA and YES. The former consists mainly of glucose, and the latter is sucrose. To investigate this, we analyzed the development and AFB_1_ biosynthesis of *A.flavus* in a toxin-favorable YES medium (20), under equal and sufficient conditions, the only variable being the carbon source (Fig. 1A). We found that the differences in growth rates in PDA and YES disappeared and that AFB_1_ synthesis did not show changes consistent with wild type (Fig. 1B). This indicates the Δ*fus3* strain loses the ability of selectively use carbon sources.

**Fig. 1.**
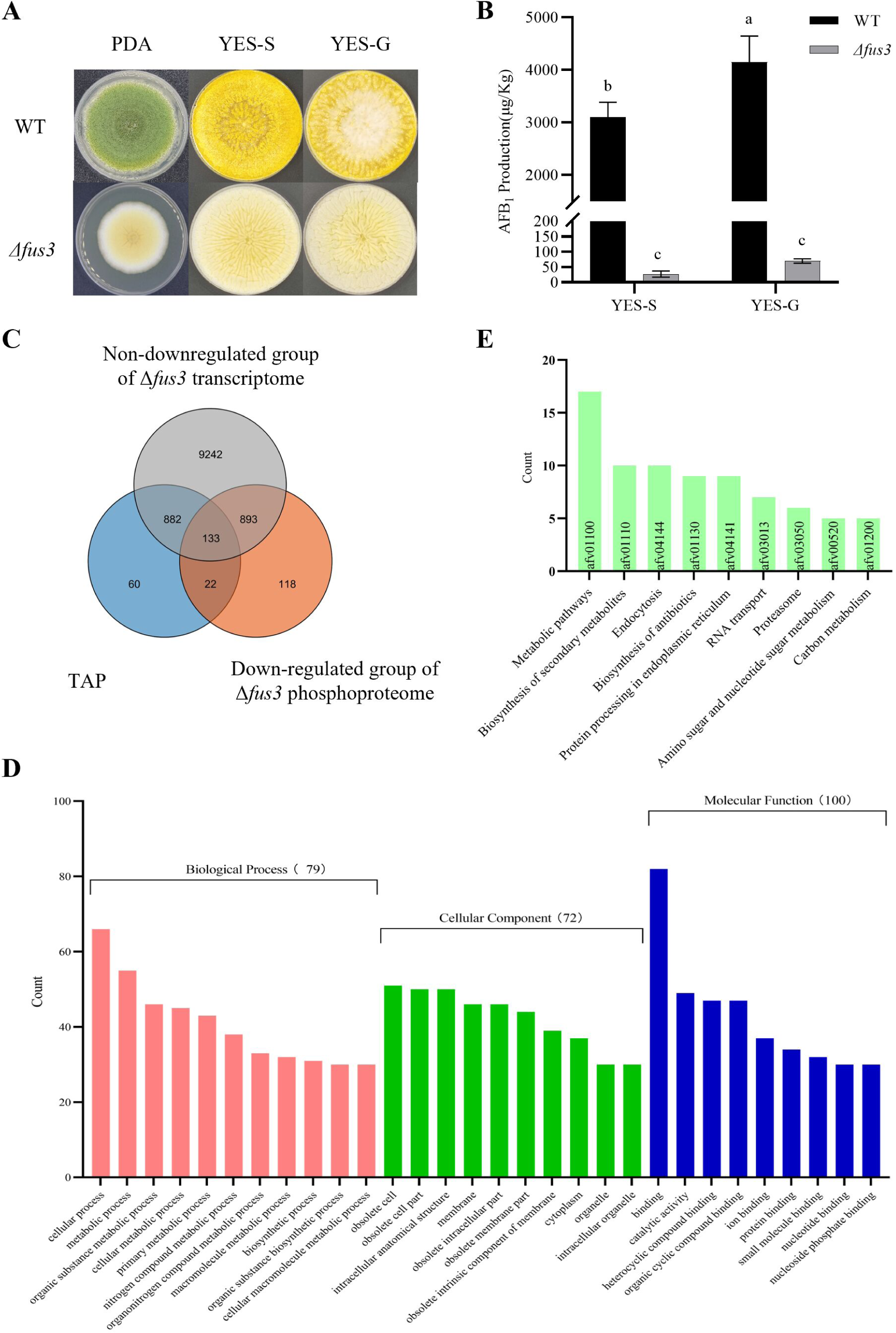
Fus3 responds to different carbon sources and screens for target proteins. (A) WT, Δ*fus3* were cultured on PDA, YES-S (Sucrose as the only source of carbon) and YES-G (Glucose as the only source of carbon) plates in 28 ℃ for 7 days. (B) the AFB_1_ production of WT, Δ*fus3* in YES-S and YES-G. The same letter indicates a non-significant difference, different letters indicate a significant difference at *p* < 0.05. Error lines indicate three biological replicates. (C) Ven diagram for cross-screening TAP, no downregulation at transcript levels and phosphorylation level down-regulation group in Δ*fus3*. (D) GO analysis of 133 screened proteins. (E) KEGG analysis of 133 screened proteins.

To find Fus3 target proteins and investigate whether they are involved in carbon metabolism, 1097 proteins were identified by the tandem affinity purification (TAP) assay in Table S1. According to the level of post-translational modification, the 133 proteins were specifically screened the phosphorylation down-regulation group (Project: IPXO003678000), the TAP group and the transcriptional level non-down-regulation group (Project: PRJNA777400) (Fig. 1C). 133 proteins that were marked by highlighting in Table S1. 55 proteins were related to the function of metabolic processes, which account for 70 % of all 79 biological process-related proteins in GO ontology analysis (Fig. 1D). 17 proteins were related to Metabolic pathway (afv01100) by KEGG analysis, which is the largest number pathway (Fig. 1E). The second largest number was the secondary metabolic pathway (afv01110) with 10 proteins. These results indicated that Fus3 is essential for carbon source selection and utilization, especially for primary metabolic pathways.

### Gal83, the β subunit of Snf1 heterotrimeric complex, interacts with Fus3 in *vivo*

According to reports (19, 21), transcription factors (TFs) might be the targets of Fus3, Sixteen proteins were selected preferentially from 133 proteins to validate interactions with Fus3 by Yeast Two-Hybrid (Y2H), involving four classes of proteins, including TFs, kinases, signal transduction proteins and molecular chaperones. Eight ORF genes out of sixteen were successfully obtained, the interaction between Gal83 and Fus3 was screened out, fortunately (Fig. 2B and Table S2). Further, Vectors crossover constructed to validate found no significant effect on the interaction relationship (Fig. 2C). The conserved domains of the Gal83 protein were analyzed by NCBI CD-search and TBtools (22) (Fig. 2A), Constructed structural domains truncation vectors and verified by Y2H found that The PRK07764 superfamily conserved domain of Gal83 did not interact with Fus3, the AMPK1-CBM conserved domain strongly interacts with Fus3, and the AMPKBI interacts with Fus3 (Fig. 2C).

**Fig. 2.**
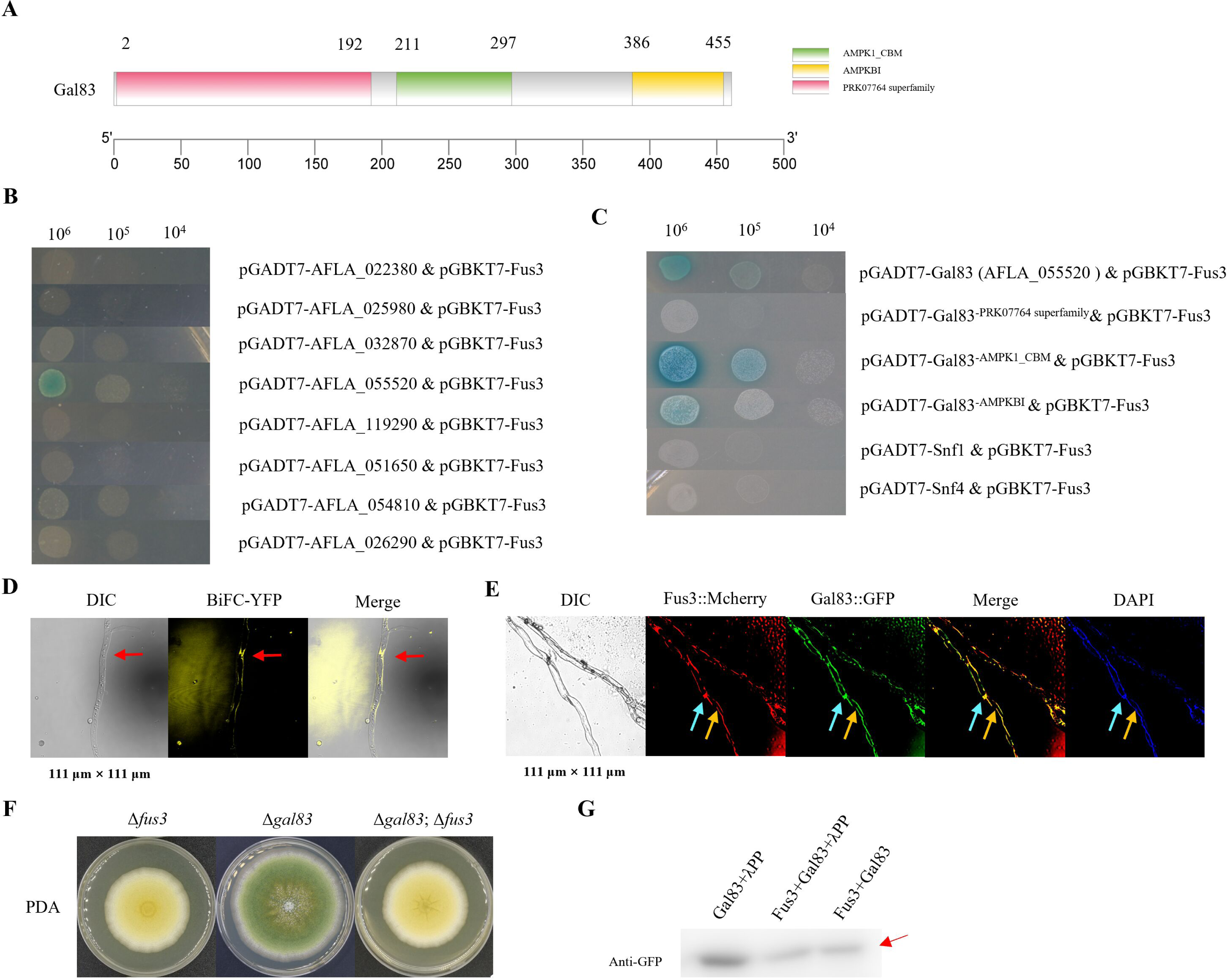
Validation of the interaction and phosphorylation relationship between Gal83 and Fus3. (A) Gal83 conserved structural domain analysis by NCBI CD-search Tool and TBtools visual graphing. (B) Screening of Fus3 target proteins from 16 proteins using yeast two-hybrid, including transcription factors, kinases, signal transduction proteins and molecular chaperones. (C) Yeast two-hybridization verified the interaction of Fus3 with each protein in the Snf1 heterotrimeric complex, including truncated sequence verification of Gal83. (D) Bimolecular Fluorescent Complimentary (BiFC) to verify the interaction between Fus3 and Gal83 in *A. flavus* in *vivo* (indicated by arrows). (E) subcellular localization of Fus3::mCherry and Gal83::GFP proteins. DAPI dyeing for membrane structures and the location of the nucleus and cytoplasm were indicated by light blue and orange arrows, respectively. (F) In *vitro* analysis of Fus3 phosphorylated Gal83, Gal83 showed a 3–5 kDA molecular weight shift (red arrow) combined with adding lambda protein phosphatase (l-PP) groups.

The BiFC-YFP (fus3::nyfp; cyfp::gal83) strain was constructed according to the bimolecular fluorescence complementation technique (BiFC), which demonstrated that Gal83 interacted with Fus3 in *vivo* under yellow fluorescence (Fig. 2D). The double fluorescent strain (fus3::mcherry; gal83::gfp) demonstrated proteins subcellular localization, with the Fus3::mCherry protein and Gal83::GFP protein in both cytoplasm and nucleus compared to the DAPI plot. The merged plot shows yellow color demonstrated the interactions, again (Fig. 2E). The above results confirmed that Fus3 interacts with Gal83 in *vivo* and *vitro* in *A. flavus*.

### Gal83 is the only target protein of Fus3 in the Snf1 heterotrimeric complex

The Snf1 heterotrimeric complex contains three subunits, α, β and γ (23). To demonstrate the interaction of Fus3 with the other proteins in the complex, firstly, we checked the TAP results found no other Snf1 complex member proteins were identified except for Gal83. We examined the genome found one each of the α-subunit protein Snf1 (AFLA_062250), the β-subunit Gal83 (AFLA_055520), and the γ-subunit Snf4 (AFLA_094030) in the Snf1 polymer. Then Y2H test shown that neither of Snf1 subunit proteins interacts with Fus3 (Fig. 2C).

The Fus3 exercises phosphorylation and Gal83 receive phosphorylation and phosphorylated α subunit Snf1, which have been demonstrated many times (1, 2, 24–27). Double gene deletion strain demonstrated that Gal83 was downstream of Fus3-Gal83 (Fig. 2F), and the protein molecular weight of Gal83 was increased after phosphorylation and western blot showed that Fus3 could phosphorylate Gal83 in *vitro* kinase assays (Fig. 2G). The screening conditions for Gal83 have demonstrated a phosphorylation transfer relationship and *gal83* deletion did not affect the phosphorylation level of Fus3 in our unpublished data. All of results supported the Gal83 was a phosphorylation and intercalation target of Fus3.

### Gal83 positively regulates development, AFB_1_ production and pathogenicity in *A. flavus*

The target protein of Fus3 is Gal83 and whether its function has similarity. The bioinformatic analysis and phenotypic identification were carried out firstly. Gal83 is conserved and the homologous protein has the highest homology in *A. flavus* and *A. oryzae* (Fig. S1A). Then we identified that obvious impairment of fungal development in *gal83* deletion strain, including the growth speed decrease (Fig. 3A and b), the conidia production significant reduction (Fig. 3A and C), the sclerotia yield severely decreased (Fig. 3A and D), and the AFB_1_ production significantly decreased (Fig. 3D). Calculating conidia production and AFB_1_ production from infested peanut and maize seeds were used to evaluate the pathogenicity of *A. flavus* (28, 29). Conidia production and AFB_1_ synthesis were significantly lower in seeds of Δ*gal83* strain infested peanut and maize than WT (Fig. 3G and H). Δ*gal83* strain exhibited the same phenotype as Δ*fus3* in YES medium with different carbon sources (Fig. 3I), also lost the ability to selectively use carbon sources (Fig. 3J). All results shown that Gal83 was synergistic with Fus3 and positively regulated mycelial growth, conidial development, sclerotia formation, AFB_1_ biosynthesis and pathogenicity.

**Fig. 3.**
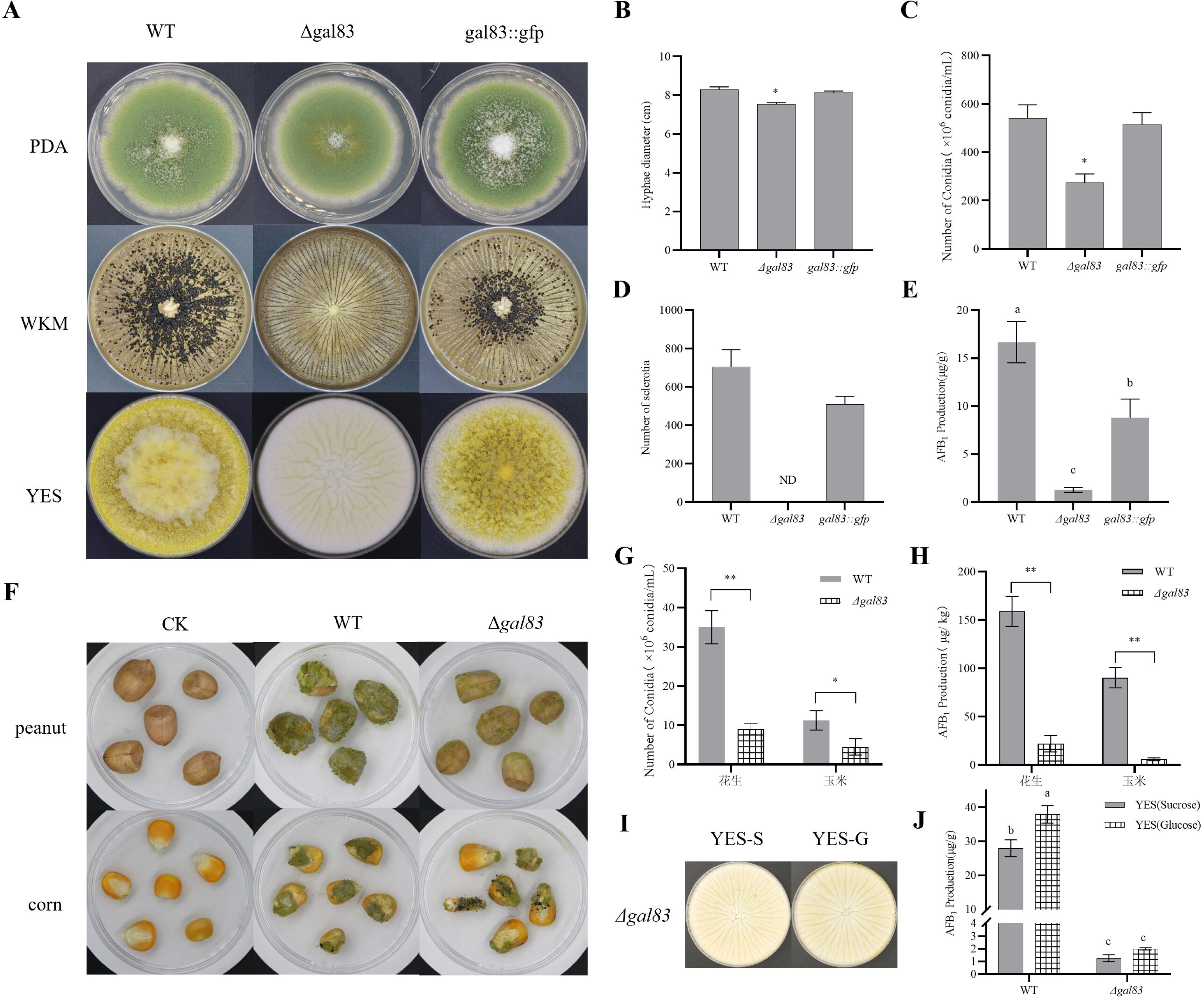
Phenotypes analysis of Δ*gal83* and complementary strains. (A) WT, Δ*gal83*, and complementary strains (Δ*gal83*; *gal83::gfp*) were cultured on PDA and YES for 7 days and WKM plates for 10 days. both PDA and WKM at 37 ℃, the YES at 28 ℃. (B) the growth rate on PDA. (C) the conidia number. (D) the sclerotia number. (E) the AFB_1_ production of WT, Δ*gal83*, and complementary strains. (F) WT and Δ*gal83* infestation of peanut and maize seeds. (G) Conidia numbers of WT and Δ*gal83* infested peanut and maize seeds. (H) AFB_1_ production of WT and Δgal83 infested peanut and maize seeds. (I) phenotype of Δ*gal83* in media with different carbon sources. (J) AFB_1_ production of Δ*gal83* in media with different carbon sources. ND, not detected. * and ** show a significant difference at P < 0.05 and P < 0.01, respectively. The same letter indicates a non-significant difference, different letters indicate a significant difference at P < 0.05. Error lines indicate three biological replicates.

### Gal83 could affect many development-related TFs at transcript levels

To unravel the mechanism of phenotype formation, we comprehensively analyzed the transcriptome of Δ*gal83* (NCBI Number) compared with Wild type. Overall, 2,854 differentially expressed genes (DEGs) were recognized in the Δ*gal83* strain (false-discovery rates [FDR] ≤ 0.05, log2FC ≥1 or ≤ -1), with 1,792 downregulated and 1,062 upregulated (Fig. S2A). The whole transcriptome DEGs results were shown in the Table S3.

We focused our analysis on important factor genes about conidia formation and developmental (Fig. 4). SteA, an important growth TF, had no significant variation at the transcriptional level. Conidial developmental genes, including conidiation-specific family protein (AFLA_044790) with -3.202 log_2_FC, conidia formation genes *abaA* and *brlA* with -4.522 and -4.135 log_2_FC, conidiation proteins *con6* and *con10* with -5.318 and -5.342 log_2_FC, and conidial hydrophobic genes *rodA* and *rodB* with -6.023 and -3.770 log_2_FC, have a significantly downregulated transcriptional level, *arb2* (-5.456 log_2_FC), *flbD* (-2.037 log_2_FC), and *wetA* (-2.540 log_2_FC) were in the same case. Only the conidial pigment biosynthesis scytalone dehydratase *arp1* (3.008 log_2_FC) has an upregulated transcriptional level. FlbA, FlbC, HymA, Con7, the conidia development-related TF, had no significant variation at the transcriptional level. The important sclerotia formation TF NsdD was no siginificant variation. Several global regulators were also analyzed, which were involved in the regulation of sclerotia formation, emergency response and secondary metabolism. Only the *velB* (-1.086 log_2_FC), which coded the protein is one of the velvet complex members with both VeA and LaeA, downregulated at transcriptional level, and the *veA*, *laeA*, *ap-1*, *atfA* were no significant variation. Overall, transcriptome analysis revealed the reasons why the growth speeds, conidia and sclerotia formation of Δ*gal83* were all defective at the transcriptional level.

**Fig. 4.**
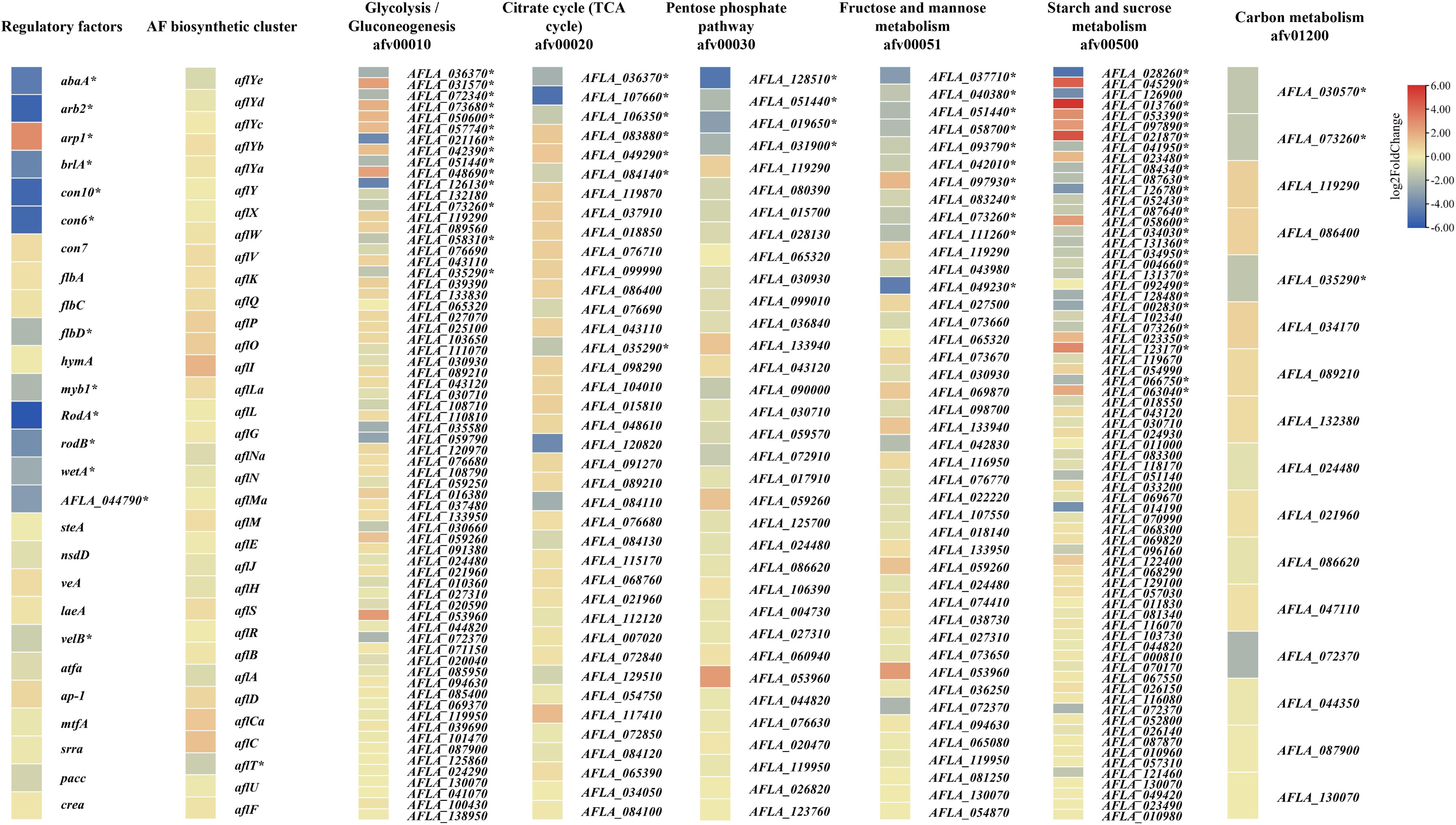
Transcriptional analysis genes of partial regulatory factors, AF synthesis gene cluster and several key metabolic pathways in Δ*gal83* vs WT. Heatmap indicates gene expression values (log_2_FoldChange), * indicates significant differences, metabolic pathway genes are categorized according to KEGG (https://www.genome.jp/kegg/). The same genes are retained at different pathways. Three biological replicates were transcriptome sequenced and agreement.

### Gal83 regulates the AFs synthetic substrates levels by regulating the key genes in primary metabolism at transcriptional level

Our previous studies have shown that Fus3 regulates Acetyl-CoA and Malonyl-CoA to modulate AFs levels in Δ*fus3* strain (1). To verify that the mechanism of AFs regulation by Gal83 was consistent with that of Fus3. Firstly, we examined and found the significantly lower levels of Acetyl-CoA and Malonyl-CoA (Fig. S2B). Then, we examined the transcript levels of the aflatoxins synthesis gene cluster, not surprisingly, there were no significant variation at the transcriptional levels (Fig. 4), RT-qPCR results were the same (Fig. S2D).

Snf1/AMPK pathway is an important that regulates metabolism (23, 24, 30). Further KEGG analyzed of the genes regulated by Gal83 in important metabolic pathways, that were categorized in many important metabolic pathways, such as the indole diterpene alkaloid biosynthesis, the arginine and proline metabolism, the starch and sucrose metabolism and the Glutathione metabolism (Fig. S2C). Several main energy metabolism pathways were checked and shown in Fig. 4 (The transcript level significantly changed genes were increased * Marker), including Glycolysis/Gluconeogenesis (afv00010), TCA cycle (afv00020), Pentose phosphate pathway (afv00030), Fructose and mannose metabolism (afv00051), Starch and sucrose metabolism (afv00500) and Carbon metabolism (afv01200). Evidently, genes downregulation occurred at the transcriptional level only in Pentose phosphate pathway (afv00030) and Carbon metabolism (afv01200) pathway. We Checked for important genes and revealed that the transcript levels of pyruvate dehydrogenase (AFLA_035290) and pyruvate decarboxylase (AFLA_126130) with -1.459 and -4.198 log_2_FC, key enzymes that catalyze the production of Acetyl-CoA from pyruvate, were both down-regulated in the Glycolysis/Gluconeogenesis, TCA cycle and Carbon metabolism pathways. The rate-limiting enzyme, 6-phosphogluconate dehydrogenase (AFLA_128510) with -4.755 log_2_FC, was the first step reaction in the Pentose phosphate pathway, which leaded to a significant down-regulation of the function of this pathway. The key enzymes in primary metabolism, including hexokinase (AFLA_073260) with -1.438 log_2_FC, phosphoenolpyruvate carboxykinase AcuF (AFLA_036370) with -2.302 log_2_FC, fructose-bisphosphate aldolase (AFLA_051440) with -1.960 log_2_FC was down-regulated at the transcriptional level. In summary, comprehensive transcriptome analysis of metabolic pathways reveal the mechanism for the downregulation of Acetyl-coA levels.

### Evidence for a complex of Fus3-Gal83-Snf1-ACCase

One of the target proteins of Snf1 is Acetyl-coA carboxylase (ACCase), which can regulate Malonyl-coA levels (23, 30, 31). We found the ACCase (AFLA_046360) was identified in both the TAP and phosphorylation downregulated results of Fus3. So, the Fus3-Gal83-Snf1-ACCase pathway was thought to be a reasonable mechanism for regulating Malonyl-coA level. To exclude direct interaction relation between Fus3 and ACCase, Y2H truncation was used for detection, because the ORF of ACCase failed to amplify from the cDNA after multiple attempts (Fig. 5A and B), and unexpected that Fus3 had strong interactions with the ACCB structural domain (Fig. 5C). We used the Global RAnge Molecular Matching (GRAMM) to show visual models of the two conjectures in Fus3-ACCase and Fus3-Gal83-Snf1-ACCase, respectively (Fig. 5D). The ACCase 3D structure of *Aspergillus* was not found, suggesting one reason why the intact ORF was not available. The model structure of *Saccharomyces cerevisiae* also suggested the possibility of intercalation, as ACCase is a conserved protein (Fig. S1B). Taken together, our results simultaneously provided evidence for direct or indirect regulation of ACCase by Fus3, which provides interesting ideas.

**Fig. 5.**
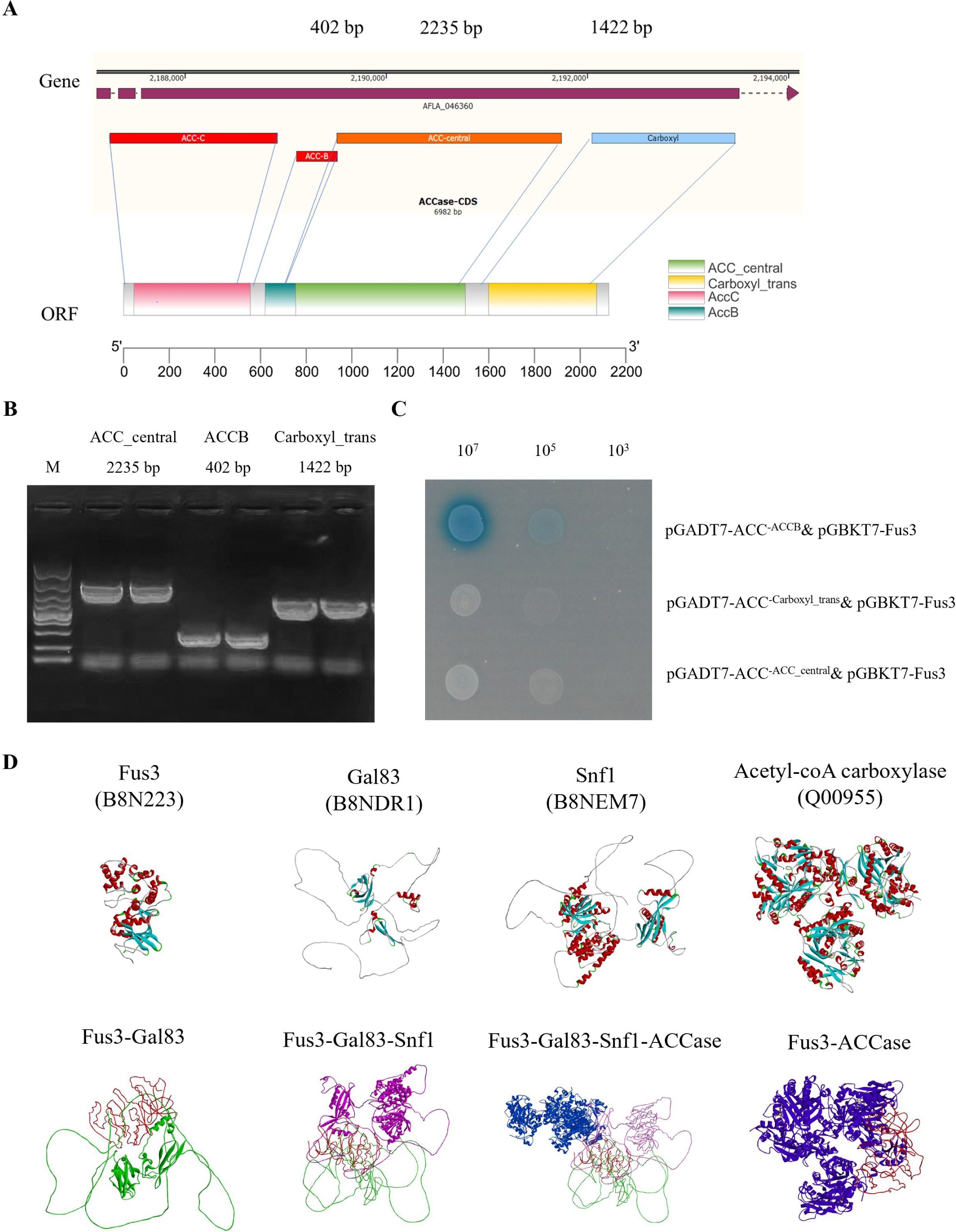
Validation and simulation of the interaction between Fus3 and Acetyl-coA carboxylase. (A) ACCase conserved structural domain analysis by NCBI CD-search Tool and TBtools visual graphing. (B) the structural domains of ACCase, PRK07764 superfamily, AMPK1-CBM and AMPKBI were amplified from cDNA. (C) Y2H verified the interactions of structural domains of ACCase with Fus3 respectively. (D) the 3D structures of Fus3, Gal83, and Snf1 proteins in *Aspergillus flavus* and ACCase protein in *Saccharomyces cerevisiae*, all data download from Uniprot (https://www.uniprot.org/). Global RAnge Molecular Matching (GRAMM) presents structural models of Fus3-Gal83, Fus3-Gal83-Snf1, Fus3-Gal83-Snf1-ACC and Fus3-ACC, demonstrating the top-rated models and visualizes.

## Discussion

Here we revealed Fus3 cross-talked to Gal83 the β subunit in Snf1/AMPK complex, a novel pathway for pheromone MAPK regulation and focused on the novel mechanism that regulates aflatoxins biosynthesis by modulating the substrates levels. Gal83 as the β subunit of the Snf1 complex that interacted strongly with Fus3 in *vivo* and *vitro* (Fig. 2C and E). The phosphorylation analysis in *vitro* and double genes deletion identified Gal83 is downstream of Fus3 and receives phosphorylation (Fig. 2F and G). Those evidence, including Acetyl-coA and Malonyl-coA levels were reduced in Δ*gal83* and Δ*fus3* strains, the aflatoxins (AFs) synthesis gene cluster was no downregulated and many key kinase genes in primary metabolisms were significant down-regulation at transcriptional level, suggested that Fus3 and Gal83 regulated AFs biosynthesis by regulating AFs substrate levels. Overall, this work uncovered new pathways in the Fus3 regulatory network and demonstrated a new theory of the Fus3-MAPK crosstalk to Snf1/AMPK pathway to regulate aflatoxin synthesis substrates. As the Fig. 6 shown, we predicted the mechanism of Fus3-MAPK crosstalk with Snf1/AMPK regulation of primery metabolism and AF synthetic substrates in *Aspergillus flavus*.

**Fig. 6.**
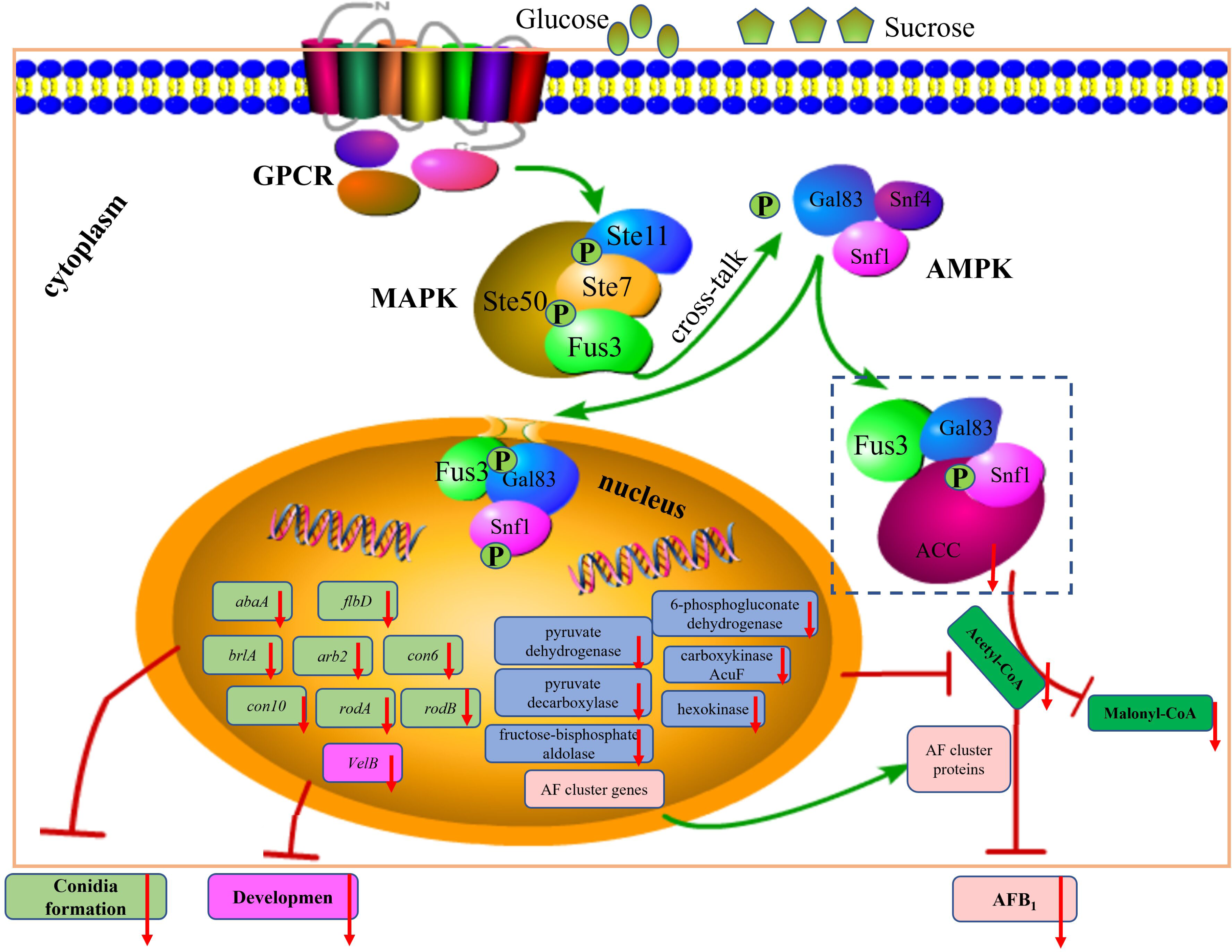
Diagram of the predicted mechanism of Fus3-MAPK crosstalk with Snf1/AMPK regulation of primary metabolism and AFs synthetic substrates in *Aspergillus flavus*. Different extracellular sugars stimulate and activate G protein-coupled receptors (GPCR), the pheromones Fus3-MAPK was activated by GPCR and carried the message. In the cytoplasm, the key kinase Fus3 was activated and interacted and phosphorylated Gal83, which as β subunit in the Snf1 heterotrimeric complex, the α subunit Snf1 protein received phosphorylation from Gal83 and then interacts with ACC to affect malonyl-coA synthesis. In the nucleus, Fus3-Gal83-Snf1 entered the nucleus and regulated the transcriptional expression of TFs, global regulators and key enzyme genes in primary metabolic pathways including the conidial formation TFs *abaA*, *brlA*, *con 6*, *con10*, *arb2*, *flbD*, *rodA* and *rodB*, the global regulator *velB*, the key enzyme genes in primary metabolic *pyruvate dehydrogenase*, *pyruvate decarboxylase*, *fructose-bisphosphate aldolase*, *6-phosphogluconate dehydrogenase*, *carboxykinase AcuF* and *hexokinase*. Finally, The MAPK-Snf1/AMPK model regulates primary metabolism, including Acetyl-coA and Malonyl-coA, the MAPK-Snf1/AMPK-ACCase (dashed box), a highly probable signaling pathway was predicted to regulate AFs biosynthesis by modulating AFs synthetic substrates.

The phosphorylation dynamics of Fus3 in *vivo* which may account for the difficulty in mining Fus3 targets. TAP is a common strategy for screening for interacting proteins in fungi (9, 15), obviously, the TAP results of the different strains differed in number and protein species. Further, the TAP results of the same *A. flavus* strain repeated more than three times in this work also differed, for example the MAPK model proteins were not identified (1) and some unnamed proteins newly appeared in the present results (Table S1). Compared to previous report results, the number of proteins in our results was greater than the Fus3-TAP result in *A. nidulans* (19), especially, the SteA/Ste12 transcription factor was not in our results. The same comparative results also appear in the results with *A. oryzae*, where our results do not have the FisA protein (15). This might be caused by the experimental time difference and the transient phosphorylation process not being easily captured. Even so, more than half of the proteins were identified repeatedly, Gal83 being one of them. This indicated that Gal83 was a stable interacting protein of Fus3. The entry of phosphorylated Fus3 and Gal83 into the nucleus (1, 19, 24, 32, 33) and the co-localization of Gal83 with it in the nucleus and cytoplasm also suggested a interaction relationship between the two proteins (Fig. 2E). Changes in Fus3 activity were more evident in the in *vitro* phosphorylation analysis, with phosphatase addition showing less significant differences in the group co-incubated with Fus3 in gal83, compared to the group without phosphatase addition (Fig. 2G). More difficult was the low biomass at relatively high expression of Fus3 and the low concentration of protein obtained, which is the reason for the shallow bands in western blot results. In addition, the low number of peptides in Fus3 was shown in the tandem affinity purification results (Table S1), this may be due to the characteristic of signal transduction proteins, which were low protein content with highly efficient.

The idea that the pheromone signaling, and glucose-sensing pathways communicate directly to coordinate cellular behavior was confirmed by Fus3-Gal83 for the first time in *A. flavus*. Sucrose non-fermenting/AMP-activated protein kinase (Snf1/AMPK) as the highly conserved serine/threonine kinases (32) and there are best known for its role in glucose limitation (34, 35). Energy metabolic processes support all life activities in cell, here, we have demonstrated that Fus3 is a specific ligand for Gal83, their stable reciprocal relationship and phosphorylation relationship had revealed the phenotype caused in *fus3* deletion. Especially the regulation mechanism of Acetyl-coA has been partially revealed by the transcriptome (Fig. 4), which is also a substrate for AFs biosynthesis. We found the β subunit to be unique in *A. flavus* and consider the role to be more important probably compared to three in yeast (36–38).

The versatility of Snf1/AMPK creates a dual effect for studies of Fus3-MAPK. As we have identified (Fig. 3), On the one hand, *gal83* deletion could explain part of Fus3 function. On the other hand, there were still many differences of function between the two proteins not been completed yet. For example, the Δ*fus3* and Δ*gal83* strains differed significantly in PDA (Fig. 1A and 3A), the acting mode of Fus3 regulated ACCase was not clear. And more importantly the mechanisms of cross-talk of different pathways have been reported, including the PKA cross-talk with Snf1 (39, 40), Snf1 cross-talk with TORC1 (41), Snf1 regulates the endoplasmic reticulum (ER) stress (42), Snf1 effects endocytosis and cell trafficking as well as cell cycle, proliferation and metabolism (43), and cross-talk between Snf1 and protein kinases involved in DNA damage (44). Overall, as the cell is the smallest unit of activity, if Fus3 is to be further resolved the impact of the unknown mechanism Snf1/AMPK must be considered. This would be a challenging and interesting study.

## Materials and Methods

### Strains, media, transformation, and cultivation of the microorganisms

The strains and plasmids used are listed in Table S4. PDA, WKM, and YES media were prepared as described previously (1, 45).

### Tandem affinity purification (TAP)

The Lapping of the *sbp*, *fus3*, *gfp* triple fragment fusion expression sequence to the *pryG* sequence by overlap PCR (FastPfu DNA Polymerase, Transgen, Beijing, China). The *pyrG-sbp::fus3::gfp* sequence was successfully constructed, then added to the homologous sequences and transformed into protoplasts of TJES 19.1 to generate the Fus3-TAP strain (46). After 72 hous incubation in YES liquid medium and protein extraction, the two kinds of Besads used for the two-step affinity purification were Anti-GFP 4FF (SMART Lifesciences, Jiangsu, China) and Streptavidin-Beads 6FF (SMART Lifesciences, Jiangsu, China) respectively. After TAP purification, the effluent proteins were harvested and then identified with LC-MS/MS.

### Yeast two-hybrid assay

The Open Reading Frame (ORF) regions of *fus3*, *gal83*, *snf1*, *snf4* and *Acc* were amplified from total cDNA, purified with a gel purification system (Magen, Beijing, China), and attached to linearized pGADT7-AD and pGBKT7-BD with a ClonExpress II One Step Cloning Kit (Vazyme Biotech Co.). The Gal83 structural domain sequences, PRK07764 superfamily, the AMPK1-CBM and the AMPKBI conserved domains were obtained to amplify from the correct *gal83* vector. The constructed plasmids were sequenced in Sangon Biotech (Shanghai, China), extracted with a TIANprep mini–Plasmid Kit (Transgene, Beijing, China), then pairwise co-transfected into the Y2HGold cells (Coolaber, Beijing, China). All of the selective medium and reagents, including SD/-Leu, SD/-Trp, SD/-Leu/-Trp, SD/-His/-leu/-Trp/-His, and X-α-gal, were purchased from Coolaber (Beijing, China). All primers used in this experiment are listed in Table S5.

### Construction of Bimolecular Fluorescence Complementation (BIFC) strain

Fus3 ORF was amplified from pGBKT7-fus3 without stop codon and fused to nyfp leading to *fus3::nyfp* fusion fragment. The *gal83* ORF was PCR-amplified from pGADT7-gal83 without stop codon followed by fusion to cyfp, which produced Fus3::nYFP and cYFP::Gal83 fusions. Primers used in this study are listed in Table S5. Null-deletion strains were constructed using a homologous recombination approach as described previously (1). The constructed fus3::nyfp with homologous sequences, was transformed into TJES19.1 to obtain the [*fus3::nyfp; pyrG*] strain. Then pyrG was deleted and the *cyfp::gal83* with homologous sequences, was transformed into [fus3::nyfp; ΔpyrG] to obtain the [*fus3::nyfp; cyfp::gal83; pyrG*] strain. To ensure sequence correctness, *fus3::nyfp* and *cyfp::gal83* sequences were constructed on the pMD18-T (D101A, Takara, Dalian, China) plasmid with ClonExpress II One Step Cloning Kit (C112-01, Vazyme Biotech Co., Nanjing, China) and sequenced. And the DNA sequences of the completed strain were also checked.

In the same way, the construction of [fus3::mCherry; gal83::gfp] strain. And the ORF of mCherry and GFP sequences were amplified from pCOM-mCherry plasmid and pCOM-GFP plasmid separately, which were from the kind gift of Dr. Li Ran and Dr. Wang dan.

### Western blotting

Both the protein extraction and the procedure of western blot were referred to Ma, et al. (1). Purification of Fus3 and Gal83 proteins using Anti-GFP 4FF (SMART Lifesciences, Jiangsu, China) and anti-mCherry Magarose Beads (ALpa-life, Shenzhen, China). Protein dephosphorylation using Lambda Protein Phosphatase (λ-PP, Beyotm, Bejing, China). Protein incubation method reference (19).

### The extraction and detection of AFB_1_ production

After 7 days of cultivation, AFB_1_ levels were determined according to Liang et al. with minor modifications (47).

### RNA extraction, reverse transcription, RT-qPCR analysis

Total RNA extraction and RT-qPCR analysis were performed according to Ren et al. (2019) and Ma et al. (2022) with minor modifications (1, 48). Important genes for AF synthesis *aflS*, *aflR*, *aflC*, *aflD*, *aflP* and *aflT* were checked in this work. The qPCR primers and ORF amplification primers are listed in Table S5.

### RNA sequencing and analysis

RNA-seq was carried out and analyzed by HTHealth (Beijing, China) according to the method described by Ma et al. (2021) (49). The gene expression changes were evaluated and the DEGs were identified with an FDR value of ≤ 0.05. Transcriptome raw data has been uploaded to NCBI (BioProject: PRJNA945182).

### Detection of acetyl-coenzyme A and malonyl-coenzyme A

Acetyl coenzyme A and malonyl coenzyme A were extracted, assayed and counted as described previously in Ma et al. (2022) (1).

### Molecular docking

PDB files of all protein 3D structures downloaded from Uniprot (https://www.uniprot.org/), Fus3 (B8N223), Gal83 (B8NDR1) and Snf1 (B8NEM7). Because the 3D model of Acetyl-coA carboxylase is not available in the genus *Aspergillus*, the file of the homologous protein in *Saccharomyces cerevisiae* was chosen (Q00955). Molecular docking tool, The Global RAnge Molecular Matching (GRAMM) operation according to instructions (50).

### Data Availability

The transcription data of Δ*gal83* and WT in this study were upload to NCBI BioProject: PRJNA945182; Both the transcriptome (Project: PRJNA777400) and phosphoproteomic data (Project: IPXO003678000) of Δ*fus3* vs WT from NCBI and the DEGs data supported by our previous research (1).

### Notes

The authors declare no competing financial interest.

## Acknowledgement

This research was funded by National Natural Science Foundation of China (31972179, 32001813), Agricultural Science and Technology Innovation Program (CAAS-ASTIP-G2022-IFST-01), Beijing Natural Science Foundation of China (6212028, 6222053), and funded by Qingdao Science and Technology Benefit the People Demonstration and Guidance Special Project (21-1-4-NY-4-NSH).

Thanks to my Professor Xing Fuguo. Thanks to my great partners: Dr. Hua Huijuan, Dr. Zhang Jun, Dr. Wang Dan, Dr. Li Ran, Lv Cong Sister, Wang Ping Sister, Ren Yaoyao, Tai Bowen.

---

**Fig. S1 The evolutionary and structural domains analyzed of Gal83 and Acetyl-CoA carboxylase proteins.** (A) and (B) are the evolutionary analyzed and structural domains of Gal83 and Acetyl-CoA carboxylase proteins respectively. Both Gal83 and Acetyl-CoA carboxylase protein sequences were obtained from NCBI, evolutionary tree analyzed by the NCBI-Blast tool and MEGA11, the Motif structural domain analyzed by the MEME tool, and the conserved structural domain analyzed by the NCBI CD-search tool.

**Fig. S2 Analysis of transcriptome results and AFs synthesis substrate levels in Δ*gal83* strain.** (A) volcano map of DEGs in transcriptome results of Δ*gal83* vs WT. (B) determination of Acetyl-coA and Malonyl-coA levels in Δ*gal83*. (C) KEGG analysis of genes with down-regulated transcript levels of Δ*gal83* vs WT. (D) RT-qPCR analysis of *aflS*, *aflR*, *aflC*, *aflD*, *aflT* transcript expression in Δ*gal83* strain. * shown a significant difference at P < 0.05. Error lines indicate three biological replicates.

